# Variation in the expression of a transmembrane protein influences cell growth in *Arabidopsis thaliana* petals by altering auxin availability

**DOI:** 10.1101/2020.01.05.895078

**Authors:** Charlotte N. Miller, Jack Dumenil, Caroline Smith, Fu Hao Lu, Neil McKenzie, Volodymyr Chapman, Joshua Ball, Mathew Box, Michael Bevan

## Abstract

**Background:** The same species of plant can exhibit highly diverse sizes and shapes of organs that are genetically determined. Defining genetic variation underlying this morphological diversity is an important objective in evolutionary studies and it also helps identify the functions of genes influencing plant growth and development. Extensive screens of mutagenised Arabidopsis populations have identified multiple genes and mechanisms affecting organ size and shape, but relatively few studies have exploited the rich diversity of natural populations to identify genes involved in growth control.

**Results:** We screened a relatively well characterised collection of *Arabidopsis thaliana* ecotypes for variation in petal size. Association analyses identified sequence and gene expression variation on chromosome 4 that made a substantial contribution to differences in petal area. Variation in expression of At4g16850 (named as *KSK*), encoding a hypothetical protein, had a substantial role on variation in organ size by influencing cell size. Over-expression of *KSK* led to larger petals with larger cells and promoted the formation of stamenoid features. The expression of auxin-responsive genes known to limit cell growth was reduced in response to *KSK* over-expression. ANT expression was also reduced in *KSK* over-expression lines, consistent with altered floral identities. Auxin availability was reduced in *KSK* over-expressing cells, consistent with changes in auxin-responsive gene expression. *KSK* may therefore influence auxin availability during petal development.

**Conclusions:** Understanding how genetic variation influences plant growth is important for both evolutionary and mechanistic studies. We used natural populations of *Arabidopsis thaliana* to identify sequence variation in a promoter region of Arabidopsis ecotypes that mediated differences in the expression of a previously uncharacterised membrane protein. This variation contributed to altered auxin availability and cell size during petal growth.

## Background

Cell proliferation and cell growth are coordinated to generate the characteristic sizes, shapes and functions of plant organs. This coordination involves multiple cellular processes, including signalling mechanisms, cell division, turgor-driven cell expansion, and cell wall and protein synthesis [1]. During the formation of determinate plant organs such as leaves and petals, cell proliferation with limited cell growth occurs at earlier stages of organ formation, followed by cell growth with limited cell proliferation occurs to increase cell size, accompanied by differentiation as the developing organ attains its final characteristic size and shape [2]. Very little is known about the spatial and temporal integration of cell proliferation and cell growth to generate the final sizes and shapes of organs and seeds, despite its fundamental and applied importance.

Many plant species display a wide range of forms due to altered sizes and shapes of organs, reflecting adaptation to their natural environments. The natural range of the annual species *Arabidopsis thaliana* extends from northern Scandinavia to Africa, and it exhibits a correspondingly diverse range of phenotypes [3–5], such that most ecotypes are phenotypically distinct. However, genetic variation underlying this phenotypic variation is still poorly described. For example, the extent to which variation in the functions of genes influencing organ size established in one widely studied ecotype influences natural variation in organ sizes in populations of *Arabidopsis thaliana* is not well understood. Also, the extent of conservation of known mechanisms influencing organ size and many other traits in natural populations is also insufficiently documented. Therefore, an increased understanding of the genetic foundations of natural variation in traits such as organ size may help to both identify mechanisms and to shed light on how natural genetic variation influences organ size and other traits.

*Arabidopsis thaliana* has adapted to diverse habitats worldwide and extensive natural variation in organ size reflects these different life histories [4]. Although variation in the shapes and sizes of different floral organs are correlated in order to maintain the reproductive functions of the flower [6], significant genetic variation influencing floral morphology traits was identified by QTL analyses of Arabidopsis Recombinant Inbred Line (RIL) populations [7, 8]. More recently, QTL analyses identified six independent loci influencing variation in petal shape and size, with variation at the *ERECTA (ER)* locus accounting for 51% of this variation [9]. Haplotype variation in 32 ecotypes at a known locus, *GA1*, was associated with variation in petal, stamen and style lengths [10]. In one of the few studies exploiting natural variation to identify *BRX* as a regulator of cell proliferation during root growth [11]. Despite these studies, there are few studies that have characterised the functions of natural variation in organ size in Arabidopsis.

Genome-wide association (GWA) mapping in Arabidopsis is increasingly used to access a wider range of natural genetic variation, to identify small-effect alleles, and to map genotype-phenotype relationships more precisely [12]. The very small size of its genomes has facilitated the re-sequencing of a large range of *Arabidopsis thaliana* ecotypes and the identification of vast numbers of SNP and small indel variants by comparison to the assembled Col-0 ecotype [13]. Within this comprehensive set of ecotypes, those from Sweden are relatively well documented [14] and have been screened for variation in over-wintering responses [15]. Initial inspection of this collection showed considerable variation in petal size and shape, therefore we conducted an association analysis of 272 sequenced Swedish ecotypes. We identified variation in the promoter of a previously undescribed gene, At4G16850, encoding a predicted 6-transmembrane (6TM) protein. Ecotypes and transgenic lines with increased At4g16850 expression had larger petals due to increased cell growth. Over-expression of At4g16850 reduced expression of several auxin-responsive genes that reduce petal cell size, and also reduced auxin availability. At4g16850 over-expression also led to the partial homeotic conversion of petals to stamenoid structures, and this was attributed to altered expression of floral organ development genes.

## Results

### Identifying a locus influencing petal area

We measured the length, maximum width and area of petals of 272 *Arabidopsis thaliana* ecotypes collected from southern and northern Sweden (Additional file 1) that were grown in controlled conditions after vernalization. Additional file 2 shows the petal phenotype data. All three petal parameters varied substantially within the sampled collection. For example, mean petal areas varied from 0.915 mm^2^ (Hov1-10) to 4.92 mm^2^ (Vår2-6), a difference of 537%. Additional file 3 shows representative petals from these ecotypes and from Död 1, an intermediate size for comparison. Figure 1A shows that mean petal area variation formed a normal distribution and was therefore suitable for association studies. GWAPP was used [16] with an Accelerated Mixed Model, incorporating information across 250,000 SNPs. This analysis identified a significant SNP association on chromosome 4 for petal area (Figure 1B). The most significantly associated SNP within this region was located at position 9471419 bp, within gene model At4G16830. The marker at this position was bi-allelic, with those ecotypes carrying an “A” allele at this locus exhibiting a ∼15% increase in petal area relative to those carrying the alternative “T” allele (Figure 1C). The extent of Linkage Disequilibrium (LD) in the region was visualised using information for all SNP markers within +/-10 kb the 9471419bp position from each ecotype and colour-coded based on the allele present. These markers, shown in chromosome order, were then sorted by phenotype values. This identified a clear block of LD (Figure 1D), with ecotypes exhibiting larger petals carrying a distinctive set of alleles from those with smaller petals. This block of LD spanned six Arabidopsis gene models, from At4g16820 to At4g16850. Sequence variation altering the activities of these genes may explain the variation in petal size observed across ecotypes. Assessment of gene annotations revealed no known regulators of petal or organ size. The effect of genetic variation within the haplotype defined by LD on petal growth was assessed in a subset of three ecotypes with small petal areas and three ecotypes with large petal areas (Figure 1E). Petal cell areas and numbers were quantified using Scanning Electron Microscopy (SEM) and Image J. A significant increase in petal abaxial epidermal cell area was observed in ecotypes carrying the increasing A allele at position 9471419 relative to ecotypes carrying the decreasing T allele (Figure 1E). Therefore, the major effect of genetic variation in the haplotype is on petal cell area.

**Figure 1.**
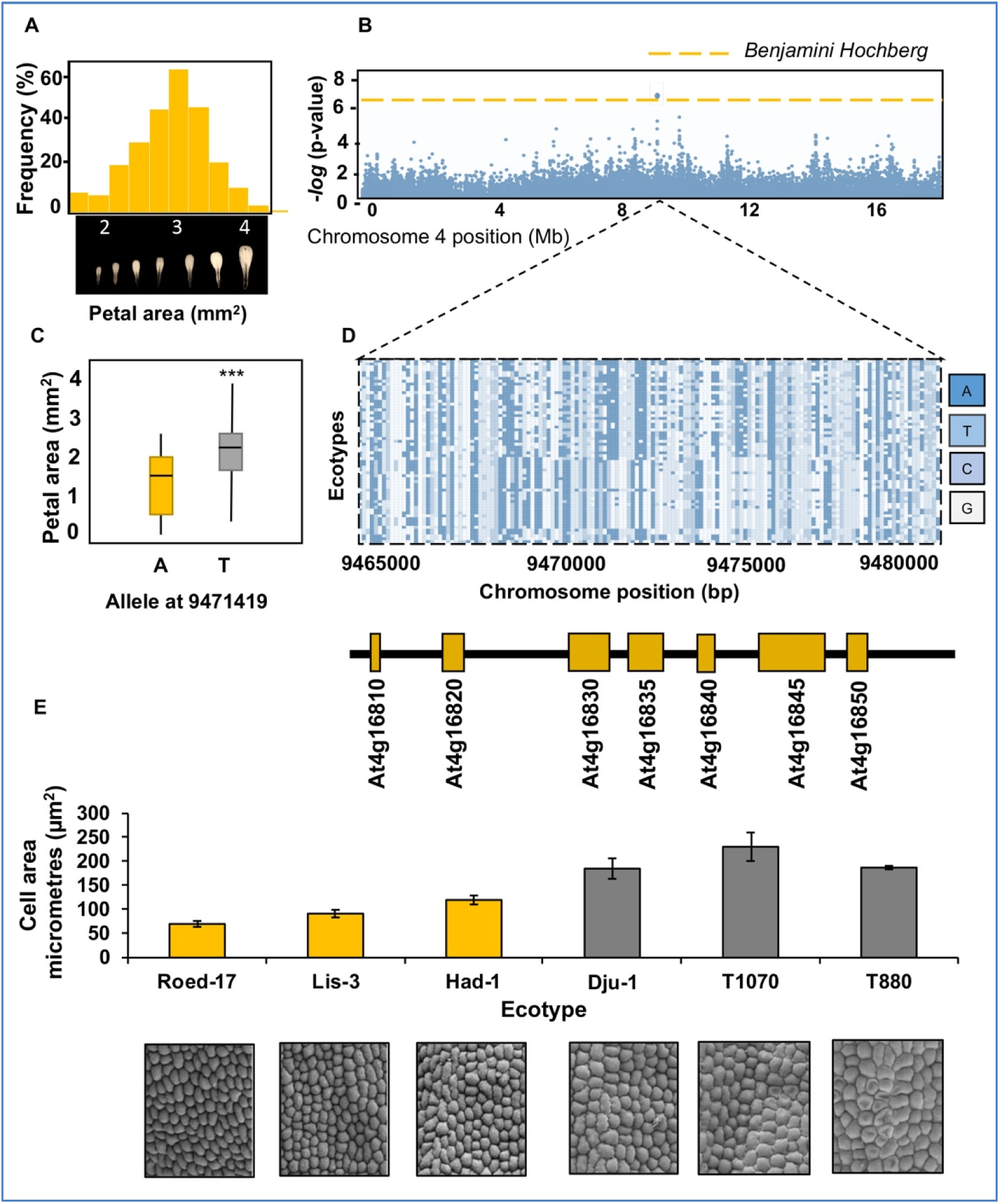
Association of petal area variation with sequence variation in Swedish ecotypes of Arabidopsis thaliana. A. Normal distribution of the petal area trait in the set of 272 ecotypes collected from Sweden (Additional file 1). B. Associations between 250,000 SNPs and petal areas determined by GWAPP using an Accelerated Mixed Model. The dotted line represents significance determined by the Benjamini-Hochberg FDR. Positions on chromosome 4 are shown in Mb, and associations as −log10 *P* values. C. Petal areas in Arabidopsis ecotypes with the A or T allele at position 9,471,419 bp on chromosome 4. *** represents *P* <0.001. D. Linkage block of haplotypes associated with large petal areas. Haplotypes extending across 15kb containing 6 gene models can be seen. The location and identity of gene models is shown in the lower panel. E. Petal abaxial cell areas of three Swedish ecotypes carrying the increasing T allele (gray bars) and three containing the reducing A allele (orange bars).

### Expression levels of At4g16850 are correlated with quantitative variation in petal size

To assess the potential role of the 6 candidate genes in regulating petal growth, petal areas were measured in available potential loss-of-function mutants in the ecotype Col-0. T-DNA insertion lines were available from stock centres for all genes found to be in high LD with associated markers with the exception of At4g16850, a small gene of unknown function. We measured the expression of these 6 candidate genes in developing floral tissues in the six ecotypes with varying petal sizes. For At4g16820 to At4g16845 no differences in petal area were seen in the T-DNA insertion lines relative to Col-0 plants (Figure 2C). Furthermore, no differential expression of these genes in developing flowers was observed between the six ecotypes with small and large petals (Figure 2B However, for At4g16850, a gene of unknown function, there was an increase in transcript levels in the three ecotypes with larger petals (P ≤ 0.001) (Figure 2B).

**Figure 2.**
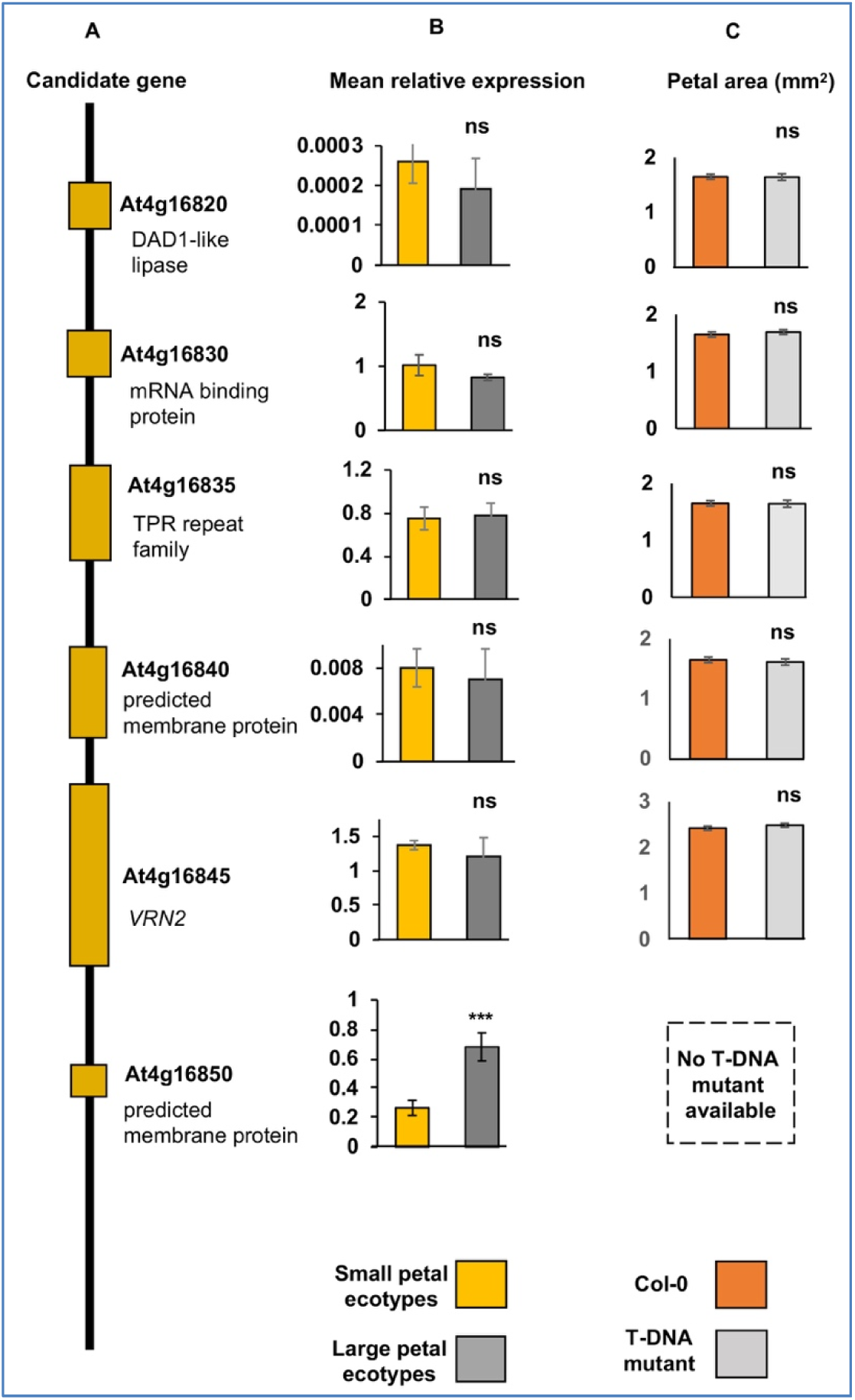
Expression levels and petal area phenotypes of candidate genes. A. Linear arrangement of predicted genes in the 15kb region of LD associated with variation for petal size. Predicted and known functions of genes in the LD block are shown. The diagram is not to scale. B. Mean gene expression levels of candidate genes in petals of three small and three large petal ecotypes. These are shown in Figure 1E. Expression of At4g16850 was significantly increased (*P*<0.001) in large petal ecotypes. Expression levels are relative to *EF1ALPHA* gene expression. Data are given as means of +/-SE (n= 3 biological replicates. *P* values were determined by Student’s *t*-test. C. Petal areas of T-DNA insertion alleles of candidate genes in ecotype Col-0. ns is not significant. No T-DNA mutant was available for At4g16850.

### Polymorphisms in the At4g16850 gene region

The relationship between increased petal areas and increased expression of At4g16850 in the selected ecotypes suggested that sequence variation between ecotypes may influence At4g16850 expression. Inspection of available genome sequence reads [13] from the larger petal ecotypes Dju-1, T1070 and T880 showed a limited range of SNP and possible small indel variation in the coding region and flanking sequences of At4g16850 compared to the Col-0 assembly. To identify any missing sequence and a wider range of promoter sequence variation, a region 2 kb upstream of the ATG initiation codon At4g16850 in three ecotypes with the decreasing A allele (Roed-17, Lis-3, Had-1) and three with the increasing T allele (Dju-1, T1070, T880) were amplified by PCR, cloned and sequenced to identify the precise location and types of sequence variation in the putative promoter regions of At4g16850. Primers were designed in conserved regions of all ecotypes. Full-length upstream regions were readily amplified from the three smaller petal ecotypes and were found to be very similar to the sequence of the Col-0 promoter (Additional file 4) carried the decreasing A allele (Figure 1C), consistent with its relatively small petal phenotype. Col-0 was therefore selected as the “small petal” reference genome due to the high level of sequence conservation between small petal ecotypes and Col-0. However, no full-length promoter amplicon could be generated from any of the large petal ecotypes. We therefore generated whole genome assemblies using Illumina sequence of un-amplified DNA templates [17] made from the three large petal ecotypes to access sequence variation in At4g16850. An ABYSS *de novo* assembly generated a large contig spanning the region upstream of the At4g16850 in line Dju-1. Comparison to the Col-0 small petal sequence identified multiple variants (Figure 3A and Additional file 4). Notably, the Col-0 and Dju-1 promoters have a common 23bp dA:dT-rich region that was extended by 30 bp in the Dju-1 promoter, making an approximately 50 bp dA:dT-rich region in Dju-1. It is likely that this dA:dT richness impeded PCR amplification of full-length upstream regions of large-petal ecotypes. There were also many other promoter polymorphisms, including another large dA/T-rich insertion in the intergenic region of Dju-1 compared to Col-0, and a deletion in Dju-1 compared to Col-0 in the 5’UTR intron (Figure 3A).

**Figure 3.**
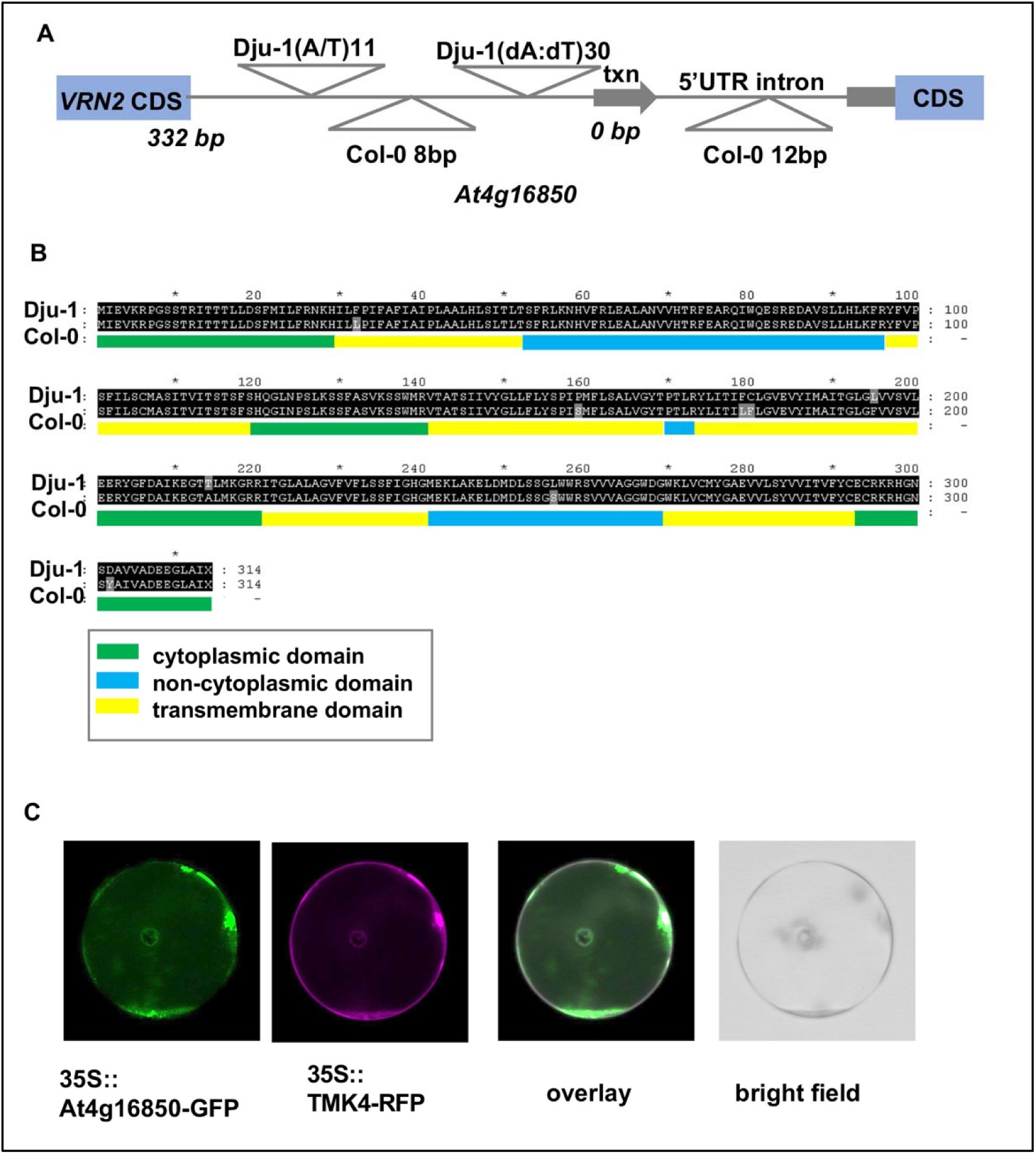
Characterisation of At4g16850. A. Sequence variation in the 322bp intergenic region between VRN2 and At4g16850 transcription start site, and in the 5’UTR, between the large petal ecotype Dju-1 and small petal genotype Col-0. Two A/T insertions in Dju-1 relative to Col-0 are shown above the line, and insertions in Col-0 relative to Dju-1 are shown below the line. Supplementary Figure 2 shows the sequence alignment of the intergenic regions. B. The coding regions of At4g16850 from the large petal ecotype Dju-1 and the small petal ecotype Col-0 were aligned to identify predicted protein sequence differences. The coding regions were analysed with InterPro to identify putative transmembrane and cytoplasmic protein domains, shown as coloured bars under the predicted coding sequence. Amino acid differences are shown as gray highlights. C. Transient expression from the 35S promoter of At4g16850 coding region fused to GFP at its C-terminus in Col-0 petal protoplasts. A known plasma-membrane protein TMK4 fused to RFP was used to reveal co-location in the plasma membrane. The white colour in the overlay reveals co-location of At4g16850-GFP and TMK4-RFP at the plasma membrane.

At4g16850 encodes a predicted 6-transmembrane domain protein with 3 non-cytoplasmic domains and 4 cytoplasmic domains (Figure 3B). Comparison of the Dju-1 and Col-0 assemblies revealed the predicted protein was highly conserved between these large- and small-petal ecotypes, with only two non-conservative amino acid changes in trans-membrane region 4 and in the C-terminal cytoplasmic domain (Figure 3B). To assess the predicted subcellular location of the protein encoded by At4g15850, its coding region was fused at its C-terminus with GFP and transiently expressed from the 35S promoter in Col-0 developing petal protoplasts, together with a known transmembrane receptor-like kinase TMK5 [18] fused to RFP. Confocal imaging showed that the At4g16850-GFP fusion protein co-localised with the RFP-tagged TMK5 plasma membrane protein (Figure 3C). The At4g16850-GFP fusion protein was also observed in cytoplasmic structures.

### Overexpression of At4G16850 increases petal size due to increased cell growth

Correlation analysis of the expression of At4G16850 across ecotypes displaying high variation for petal area established that its differential expression explained 76% of the variation in petal size in the analysed ecotypes (Figure 4A). To establish whether this variation in At4g16850 caused petal size variation, the coding region of At4g16850 from Col-0 was expressed from the constitutive 35S promoter in transgenic Arabidopsis Col-0 plants. Col-0 has relatively small petals and inherits the decreasing allele in the associated haplotype that segregates with low At4G16850 expression.

**Figure 4.**
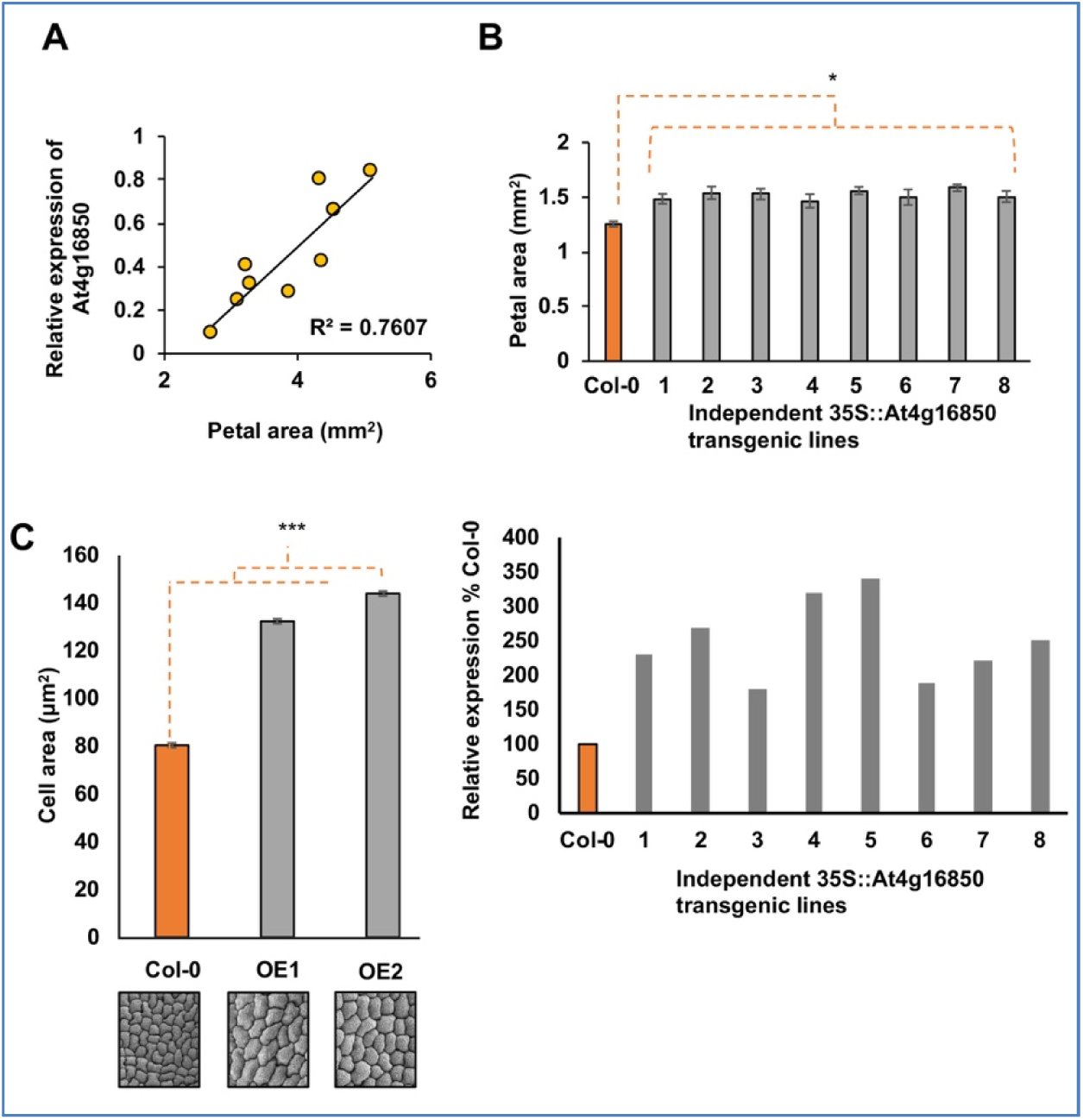
Over-expression of At4g16850 leads to larger petal cells. A. Correlation of At4g16850 gene expression and petal areas in ecotypes exhibiting a wide range of petal areas. B. Petal areas in transgenic Col-0 lines overexpressing At4g16850 from the 35S promoter. Eight independent transgenic lines with elevated At4g16850 transcript levels in petals were selected and petal areas measured (n=50 petals, *P<0.05). The lower panel shows relative expression levels of At4616850 in the transgenic lines compared to Col-0. C. Cell areas on the abaxial side of petals from two selected over-expressing lines described in panel B above are increased (***P<0.001, n=50 cells).

Comparison of petal areas in transgenic lines and untransformed Col-0 plants revealed that transgenic plants overexpressing At4G16850 (Figure 4B lower panel) exhibited significantly increased petal size relative to WT plants (P ≤ 0.01) (Figure 4B upper panel). No other phenotypic differences between Col-0 and the transgenic lines were observed. Therefore, increased expression of At4g16850 leads to increased petal area, showing that variation in At4g16850 expression among the ecotypes can directly influence petal area. In the ecotypes exhibiting increased At4g16850 expression, cell areas were increased and accounted for larger petal areas (Figure 1E). Petal cell areas were also increased in transgenic lines overexpressing At4g16850 (Figure 4C). Taken together, these results show that increased expression of At4g16850 promotes cell growth in Arabidopsis petals. To take account of this information about a previously unknown gene in *Arabidopsis thaliana* we named the gene *KSK* (<u>K</u>ronblad<u>S</u>torle<u>K</u>, Swedish for petal size).

### Increased expression of *KSK* reduces expression of genes that limit petal cell growth

Previous studies have identified several genes that influence petal cell growth in Arabidopsis. *BPEp* [19] and *ARF8* function together [20] to limit petal cell growth, and *FRL1* [21] also represses petal cell growth. The expression of these genes in developing petals of three transgenic Col-0 lines over-expressing At4g16850 and in untransformed Col-0 was measured using Q-RT-PCR to assess whether *KSK* may influence petal cell growth through these genes. Although only one transgenic line showed significant reduction in *BPE* expression in petals (Figure 5A), consistent reductions in *ARF8* and *FRL1* expression in developing transgenic petals was seen (Figures 5B, C) compared to Col-0. This suggested that *KSK* may promote petal cell growth by reducing the expression of these petal cell growth genes. *AGAMOUS* reduces BPEp expression [19] and the *ag1* loss of function mutant has larger petals [22], consistent with the larger cell and petal sizes in BPEp loss of function mutants. Although we did not observe consistent reduction of *BPEp* expression in all *KSK* overexpressing transgenic lines, we tested whether *AG* influences *KSK* expression. *KSK* expression was doubled in the *ag1* loss-of-function mutant (Figure 5D) consistent with a model in which *AG* represses *KSK* expression, leading to increased *ARF8* and *FRL1* expression and corresponding reduced petal cell size and overall petal size.

**Figure 5.**
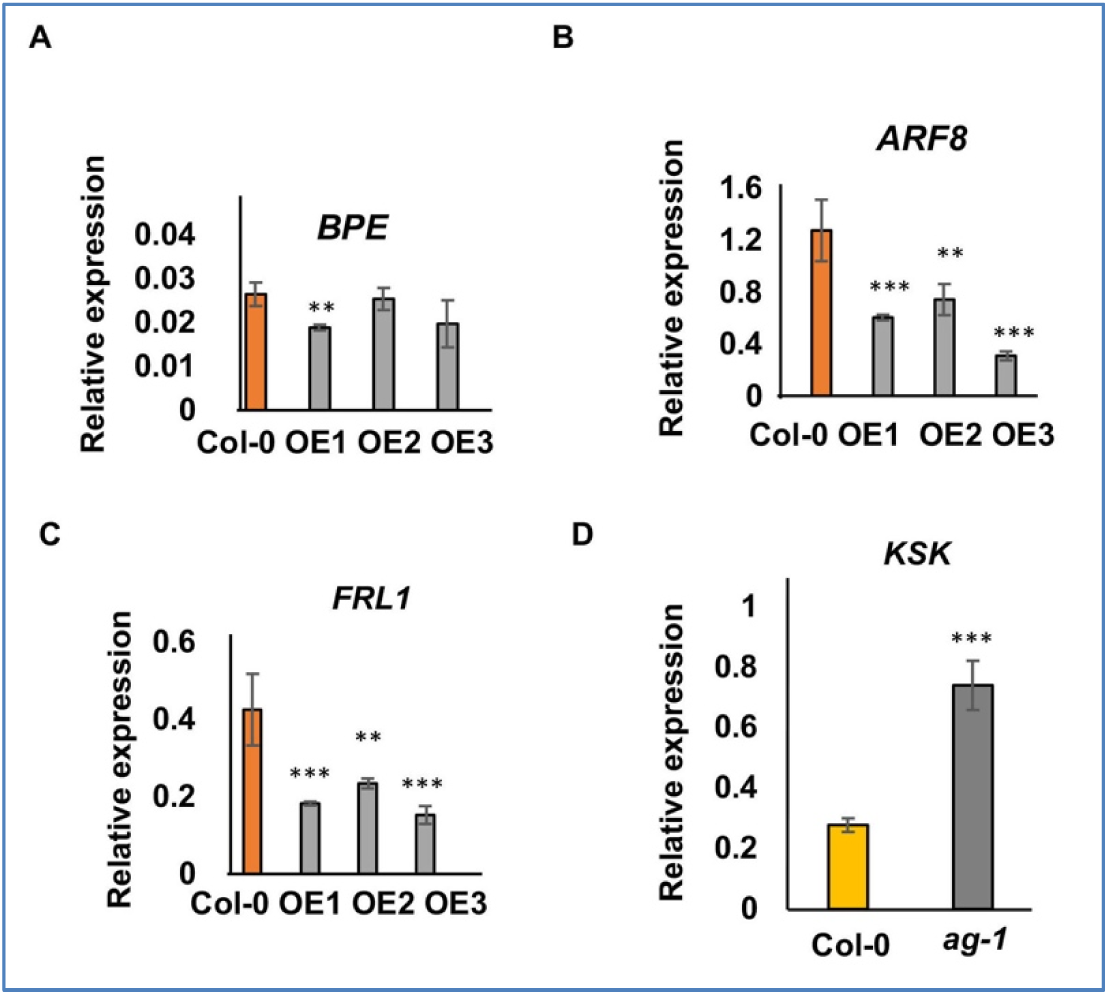
Reduced expression of petal cell growth genes in *KSK* over-expressing lines. Expression levels of *BPE* (A), *ARF8* (B) and *FRL1* (C) in developing floral tissues of three *35SS::KSK* overexpressing lines (OE 1,2,3). ** *P*< 0.01; *** *P*<0.001. Expression levels are relative to *EF1ALPHA* gene expression. Data are given as means of +/-SE (n= 3 biological replicates. *P* values were determined by Student’s *t*-test. D. Expression of *KSK* in developing petals of Col-0 and the *agamous 1* mutant. Expression levels are relative to *EF1ALPHA* gene expression. Data are given as means of +/-SE (n= 3 biological replicates. *P* values were determined by Student’s *t*-test.

### Overexpression of *KSK* leads to partial homeotic conversion of petals to stamenoid structures

In addition to observing a significant increase in petal cell growth in the *35S::KSK* over-expressing lines, we also observed partial organ identity changes in ∼10% of flowers from all eight *35S::KSK* transgenic plants. These flowers, in addition to the expected four petals in the second whorl, carried a 5th petal-like structure. This developed in the outer margin of the second whorl and displayed stamenoid features such as a partial pollen sac (Figure 6A) and stomata, a cell type not observed in Col-0 petals (Figure 6B). The presence of stomata on petal epidermal surfaces has also been seen in *ant* mutants deficient in the transcription factor *AINTEGUMENTA* (Krizek 2000). Using qRT-PCR, we assessed *ANT* expression in developing petals of the three *35S::KSK* over-expressing lines. A significant decrease in *ANT* expression was observed in petals overexpressing *KSK* (Figure 6C). This suggests that *KSK* expression levels contribute to determining floral organ identity in a pathway involving *ANT*.

**Figure 6.**
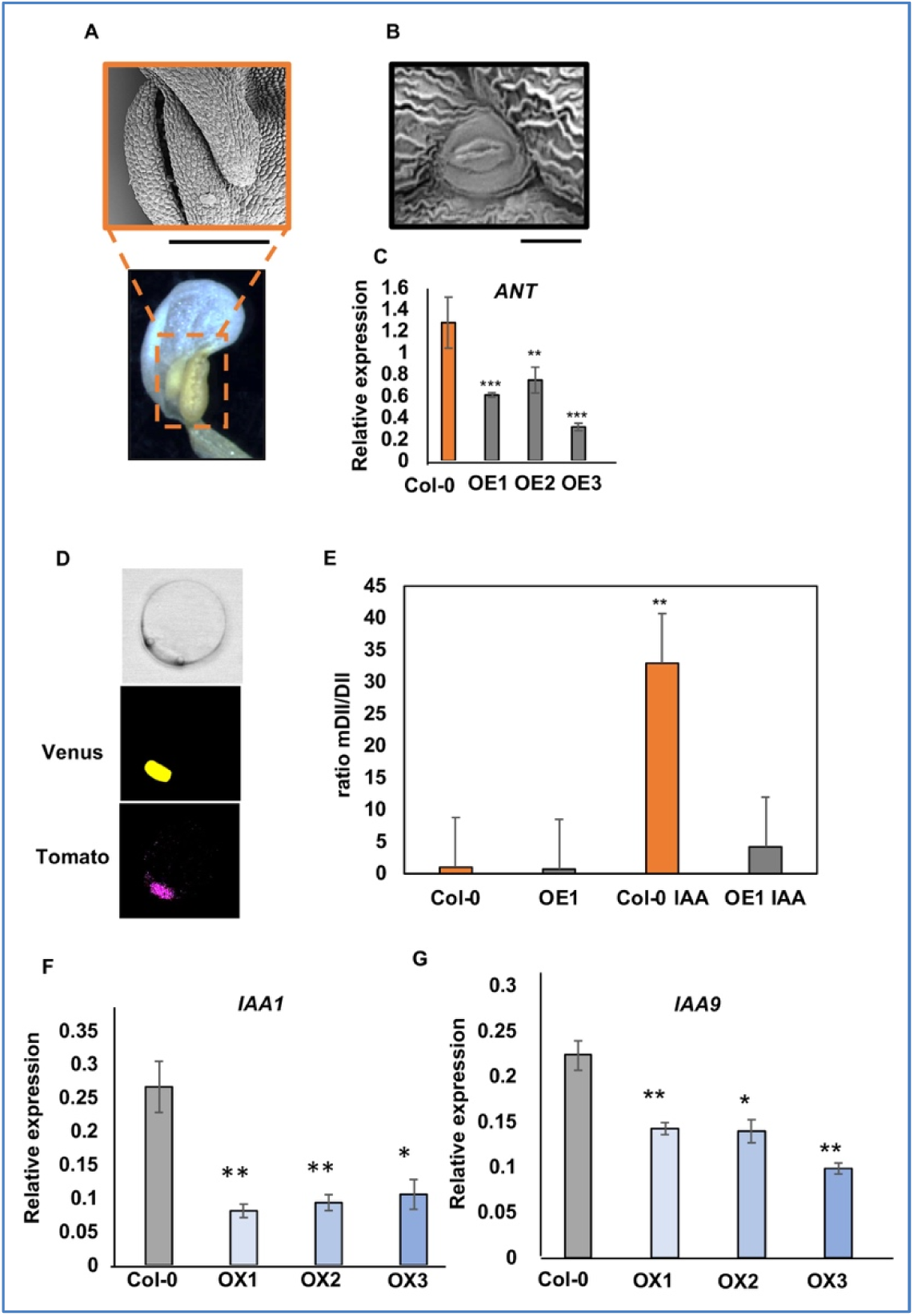
Over-expression of *KSK* leads to partial homeotic conversion of petals to stamenoid structures and leads to reduced auxin responses. A. SEM (upper) and bright field (lower) image of a petal from a *35S::KSK* transgenic line showing stamenoid features such as a pollen sac. The scale bar represents 100 μm. B. SEM of petal epidermal cells from a *35S::KSK* transgenic line showing a stomata. The scale bar represents 10 μm. C. Relative expression of *ANT* in Col-0 and *35S::KSK* petals. Expression levels are relative to *EF1ALPHA* gene expression. Data are given as means of +/-SE (n= 3 biological replicates. *P* values were determined by Student’s *t*-test. ** *P*< 0.01; *** *P*<0.001. D. Nuclear localisation of DII-Venus (middle panel) and mDII-Tomato (lower panel) from transiently expressed R2D2 in Col-0 petal protoplasts. The upper panel is the imaged protoplast in bright field. E. Ratios of mDII-Tomato to DII-Venus transiently expressed in Col-O or *35S::KSK* petal protoplasts. Protoplasts (from line OE1) were treated with 0 nM or 1000 nM IAA for 1-2 hrs before imaging. The increased ratio of mDII/DII shows reduced levels of DII-Venus in response to elevated auxin levels. F and G. Reduced expression of the auxin-responsive genes *IAA1* (Panel F) and *IAA9* (Panel G) in Col-0 and three transgenic lines over-expressing *KSK*. Expression levels are relative to *EF1ALPHA* gene expression. Data are given as means of +/-SE (n= 3 biological replicates. *P* values were determined by Student’s *t*-test.

### Reduced auxin responses in lines over-expressing *KSK*

One common feature of *ARF8* and *FRL1* expression, which was suppressed by over-expression of *KSK*, is that the expression of both genes increases in response to auxin [20, 23, 24]. We therefore tested auxin responses in petals of wt Col-0 and *35S::KSK* over-expressing lines. The DII-n3Venus reporter protein is a fusion of the auxin-dependent DII degron to a nuclear localised Venus reporter coding region. Auxin levels are detected by reduced Venus fluorescence relative to a mutant form, mDII-ntdTomato, which is not degraded in the presence of auxin [25]. This dual auxin reporter, called R2D2, is suitable for single cell assays as different transformation efficiencies can be accounted for by the relative fluorescence of nuclear-localised Venus and Tomato fluorescent proteins. Protoplasts were isolated from developing petals from Col-0 and a *35S::KSK* transgenic line over-expressing *KSK* and transfected with the R2D2 plasmid. After 16 hrs to allow for protein expression, protoplasts were treated with either 0 nM or 1000 nM IAA. Protoplasts were imaged between 1-2 hr after IAA treatment. Figure 6D shows the nuclear localisation of both fluorophores and Figure 6E shows relative fluorescence of mDII-Tomato/DII-Venus measured in the nuclei of transfected petal protoplasts. The increased ratio in Col-0 protoplasts shows a reduction in DII-Venus protein compared to mDII-Tomato in response to auxin, demonstrating that transiently-expressed protoplasts respond to added auxin similarly to stable transgenic plants [25]. In contrast, in *35S::KSK* transgenic protoplasts, the ratio of Venus to Tomato fluorescence was not decreased to the same extent as Col-0. This indicated that auxin responses may be reduced in this transgenic line. This interpretation was tested by measuring the expression levels of two auxin-responsive genes (*IAA1* and *IAA9*) in seedlings of Col-0 and *KSK* over-expressing lines. Their expression was reduced (Figures 6F,G), supporting the interpretation that auxin responses are decreased in *KSK* over-expressing lines.

## Discussion

Approximately 7,000 *Arabidopsis thaliana* ecotypes have been systematically collected and 1,000 of these have been sequenced to access a wide range of genomic diversity [13]. We used genome-wide association analyses [26] on a set of sequenced ecotypes collected from Sweden [14] to identify a major source of sequence variation influencing petal size. Variation in the expression of a gene encoding a hypothetical plasma membrane protein, which we call *KSK*, made a major contribution to variation in petal size in this set of ecotypes. Auxin responses as measured by the R2D2 reporter were reduced in *KSK* over-expression lines, suggesting that the KSK membrane protein may directly or indirectly influence auxin responses or levels in developing petals.

*KSK* was predicted to encode a transmembrane protein with 6 short helical domains spanning the membrane, 3 non-cytoplasmic domains and 4 cytoplasmic domains in an N-in conformation (Figure 3B). A KSK-GFP fusion protein was co-localized with a plasma membrane protein (Figure 3C). The KSK protein sequence is reasonably highly conserved among several groups of plants, and it has no close family members in Arabidopsis, with only very partial protein alignments to At1g31130 detected by reciprocal BLASTP searches. At1g31130 is a 321aa predicted 6TM protein located in Golgi, endosomes and the plasma membrane. The 6TM protein structure is predicted to be present in many Arabidopsis proteins with diverse functions, including aquaporins, voltage-gated ion superfamily transporters, and mitochondrial carrier proteins [27]. The MIND1 database of membrane protein interactions [28] identified an interaction between *KSK* and At1g07860, encoding a predicted Receptor-Like Cytoplasmic Kinase VII (RLCK VII) family member. Members of this family function in pathogen-triggered immunity and growth pathways [29] and include BIK1, which is membrane anchored *via* N-myristoylation [30]. In *KSK* over-expressing lines, auxin responses were reduced as detected by transient expression of the R2D2 auxin reporter gene (Figure 6E). This may be due to reduced auxin responses, synthesis, or altered transport. The membrane localisation of KSK-GFP suggests a potential influence on auxin transport. However, the predicted 6TM transmembrane organisation of KSK in membranes is different from that of all known auxin uptake and efflux plasma membrane- and tonoplast-located auxin transporters [31].

Multiple promoter polymorphisms were identified between the small petal ecotype Col-O and the large petal ecotype Dju-1 (Figure 3A and Additional Figure 2) after re-sequencing and assembling Dju-1. In contrast, the protein coding regions of these two ecotypes had only two non-conserved amino acid changes (Figure 3B); one was in a predicted non-cytoplasmic domain and the other in the predicted C-terminal non-cytoplasmic domain. While these may influence *KSK* function in different ecotypes, the phenocopying of the big petal phenotype of Dju-1 by over-expressing the Col-0 coding region indicated that promoter variation is most likely causal for the large petal phenotype in the screened ecotypes. An interesting feature of the Dju-1 promoter region is the accumulation of expanded dA:dT-rich tracts of up to 30 bp. Polymorphisms of this length are very common in Arabidopsis genome assemblies [32], and are over-represented in many eukaryotic genomes, where they may be generated by replication slippage [33]. dA:dT tracts in promoters have a well-established role in regulating gene expression by forming part of scaffold attachment regions (SARS) and by introducing curvature in DNA that influences transcription factor and nucleosome access.

*KSK* over-expression led to reduced expression of *ARF8* and *FRL1*, two genes that exert a specific negative effect on petal cell size (Figures 5A,B,C). *FRL1* encodes a sterol methyltransferase that influences endoreduplication [34]. ARF8 is an auxin-responsive transcription factor that forms a transcription complex with the bHLH transcription factor BPEp [20]. BPEp also restricts cell expansion specifically in petals. BPEp is highly expressed during the later stages of petal development, while ARF8 is ubiquitously expressed, but more highly expressed during the later stages of petal development. The expression of both *FRL1* and *ARF8* is increased in response to auxin [23, 24, 35], and as auxin responses are reduced in *KSK* over-expressing lines (Figure 6E), it is possible that *KSK* over-expression may reduce expression of these auxin-responsive negative regulators of petal cell size, leading to increased petal cell size and overall increases in petal area. The down-regulation of *ANT* expression in *KSK* over-expression lines (Figure 4C) is consistent with reduced auxin responses, as auxin increases *ANT* gene expression [36]. *ANT* encodes an AP2/ERF transcription factor that influences several stages of floral development, including specification of floral organ identity [37]. Petals in *KSK* over-expressing lines often exhibited a partial conversion to stamenoid features, and also had stomata, a cell type not normally found in petals (Figures 6A, B). In *ant* mutants, petal cell identity was also altered to form stomata [22], supporting the interpretation that *KSK* modulates *ANT* expression, perhaps by altered auxin responsiveness, leading to partial homoeotic conversion of petals to stamenoid features. Such conversion is consistent with another function of *ANT* in restricting, together with *AP2*, *AGAMOUS (AG)* expression to the second whorl. AG activates expression of *SPOROCYTELESS/NOZZLE (SPL/NZZ)*, a transcription factor that promotes microsporogenesis [38]. A lessening of this restriction of *AG* expression could therefore conceivably lead to stamenoid features developing on *KSK* over-expressing petals.

Floral morphology plays a central role in plant fitness, for example by attracting specific pollinators. In *Brassica napus* crops, reduced petal size is an important trait as it increases light penetration through dense canopies. In wild populations of *A. thaliana*, the adaptive significance, if any, of varying petal areas is not well understood. Although *Arabidopsis thaliana* is primarily self-pollinating, some populations exhibit elevated outcrossing, especially in species-rich rural environments [39]. Diverse insects visit Arabidopsis flowers and are potential pollinators [40], therefore it is reasonable to speculate that genetic variation in petal size may have an adaptive role in securing outcrossing in Swedish populations. The three lines selected with large petals came from different locations in southern Sweden (Additional Table 1), all of which harbour ecotypes with varying sized petals. Natural variation in petal size in *A. thaliana* may therefore contribute to diversifying outcrossing opportunities by attracting different types of insect visitors.

## Methods

### Plant material and phenotyping

Arabidopsis GWAS ecotypes and all transgenic plants and T-DNA mutants were grown on soil in a growth chamber with 16/8 hr day/night at 22°C after 48 hours stratification at 5°C. To quantify petal lengths, widths and areas, 10 petals were dissected from the first set of completely open flowers from 3 biological replicates per genotype to minimise any developmental differences. Petals were mounted on black cardboard and laminated to protect the petals. Petals were scanned at 200dpi resolution and petal length, maximum width and areas measured using Image J.

### GWAS

GWAS was carried out using the open-source GWAS software, GWAPP: https://gwas.gmi.oeaw.ac.at/ with the 250K SNP dataset. GWAS was performed using the AMM function.

### DNA constructs

The *p35S::3xHA-At4g16850* transgenic line was generated by cloning *At4g16850* cDNA into the pENTR TOPO-D vector (Thermofisher, UK) using the primers described in Additional Table 2. LR Clonase mix II (Thermofisher, UK) was used to transfer the *At4g16850* CDS into the 35S PB7HA binary vector. The pAT4g16850::AT4G16850-GFP transgenic line was generated by cloning the promoter and coding region from genomic DNA into the pENTR TOPO-D vector. LR clonase was then used to transfer the target sequence into the pEARLYGate 103 vector. All constructs were sequenced before use. The *p35S::3XHA-At4g16850* construct was transformed into *Agrobacterium tumefaciens* strain GV3101, and Arabidopsis *Col-0* plants were transformed using the floral dip method [41].

### Genotyping Arabidopsis T-DNA lines

Sequence indexed T-DNA insertion lines, obtained from The Nottingham Arabidopsis Stock Centre (NASC), were genotyped using gene-specific primers designed using the primer design tool http://signal.salk.edu/tdnaprimers.2.html: Primer sequences are in Supplementary Table 2. Genotyping was carried out using TAKARA EX taq (Takara Bio, USA) according to the manufacturer’s instructions.

### cDNA synthesis, PCR and Genome Sequencing

RNA was extracted from developing floral buds using the SPECTRUM Total Plant RNA kit (Sigma, UK). 1ug of RNA was treated with RQ1 RNase-Free DNase (Promega, USA) and cDNA synthesis was carried out using the GoScript Reverse Transcription system (Promega, USA) using OligoDT. All protocols were carried out using manufacturers’ guidelines. cDNA samples were diluted 1:10 in water before use. Q-RT-PCR was carried out using SYBR green real time PCR mastermix (Thermofisher) and performed using Lightcycler 480 (Roche, Switzerland). Primer sequences used for q-RT-PCR are in Additional file 5. Primer efficiencies and relative expression calculations were performed according to methods described by [42]. All q-RT-PCR assays were repeated at least twice. All PCR reactions were carried out using Phusion High Fidelity DNA polymerase (New England BioLabs) according to manufacturer’s instructions. Capillary sequencing was carried out by GATC Biotech (Germany). For whole genome assembly of ecotypes Dja-1, TBA_01 and TI_070, high MW DNA was prepared and PCR-free indexed Illumina libraries prepared as described [43]. After QC approximately 50m 150bp paired-end reads were generated (Novagene, Hong Kong) for each library. Cleaned reads were assembled using ABySS v1.3.6 [44] with a *k*-mer of 75. Genome assemblies were aligned with the genomic region of At4g16850 using MUSCLE v3.8.31 [45]. Assemblies of the three ecotypes are available at ENA (PREJB28030).

### Scanning Electron Microscopy

Petals were dissected, fixed and critical point dried before SEM imaging. Chemical fixation was in 2.5% glutaraldehyde in 0.05 M sodium cacodylate, pH 7.4. Vacuum infiltration was carried out until the petals sank before leaving overnight in fixative at 4° C. After rinsing in buffer twice and then water twice for 15min each, petals were dehydrated through an ethanol series for 30 min each in 30%, 50%, 70%, 90%, 100%, and 100% dry ethanol, then critical point dried using a Leica EM CPD300 system (Leica Microsystems Ltd, Milton Keynes, UK) according to the manufacturer’s instructions. Dried samples were mounted on the surface of an aluminium pin stub using double-sided adhesive carbon discs (Agar Scientific Ltd, Stansted, Essex). The stubs were then sputter coated with approximately 15nm gold in a high-resolution sputter coater (Agar Scientific Ltd) and transferred to a Zeiss Supra 55 VP FEG scanning electron microscope (Zeiss SMT, Germany). The samples were viewed at 3kV and digital TIFF files were stored.

### Transient expression in Arabidopsis protoplasts

Transient expression assays were carried out using protoplasts isolated from *Arabidopsis* Col-0 developing petals [46]. Protoplasts were transformed with 5ug plasmid DNA (purified using the Qiagen Plasmid Maxi Kit (Qiagen)). After an overnight incubation at 20°C, transfected protoplasts were harvested and imaged using confocal microscopy. The R2D2 plasmid (Addgene 61629 pGreenIIM RPS5A-mDII-ntdTomato/RPS5A-DII-n3Venus) was transfected into petal protoplasts isolated from Col-0 or *35S::KSK* plants. After overnight incubation, protoplasts were treated with 0 or 1000nM IAA for 1-2 hrs before imaging. A Leica SP5 set up for photon counting at 12 bit resolution was used for imaging transfected protoplasts. Gain was set to 50% for Tomato fluorescence and to 10% for Venus fluorescence. Venus was excited at 514nm and detected at 524-540nm, and Tomato was excited at 561nm and detected at 571-630nm. ImageJ was used to calculate the mean gray value of fluorescence within nuclei.

## Supplementary Information

Additional file 1: Table S1. List of Arabidopsis thaliana ecotypes used in the GWAS analysis.

Additional file 2: Table S2. Petal phenotype data.

Additional file 3: Figure S1. Representative petals from the Swedish ecotypes.

Additional file 4: Figure S2. Promoter alignments of the Dju-1 and Col-0 *KSK* genes.

Additional file 5: Table S3. Primer sequences used in this study.

## Acknowledgements

We are grateful to members of Caroline Dean’s group at the John Innes Centre for assistance with the Swedish ecotypes, and to staff from the John Innes Centre Horticultural Services for expert plant maintenance.

## Author Contributions

CNM, JD, MB and MWB conceived the experiments, MWB supervised the work, CNM, JD, F-HL, CS, NMcK, VC and JB carried out the work, CNM and MWB wrote the paper.

## Data Availability

Sequence reads of the *A. thaliana* ecotypes Dju-1, TBA_01 and TI-070 is available at ENA (PRJEB28030).

