## Supplementary material for "Variation in the expression of a transmembrane protein influences cell growth in *Arabidopsis thaliana* petals by altering auxin availability": Table S1

| **AccessionName** | **AccessionID** |
| --- | --- |
| T910 | 6143 |
| Ådal 3 | 9323 |
| Öde 2 | 9434 |
| Öde 3 | 9435 |
| Ale1-2 | 5829 |
| AleA 1 | 9325 |
| Aledal-11-63 | 1163 |
| Aledal-1-34 | 1153 |
| Aledal-14-73 | 1166 |
| Aledal-17-82 | 1169 |
| Aledal-6-49 | 1158 |
| Ale-Ster-41-1 | 991 |
| Ale-Ster-44-4 | 992 |
| Ale-Ster-50-11 | 996 |
| Ale-Ster-56-14 | 997 |
| Ale-Ster-57-16 | 998 |
| Ale-Ster-59-18 | 999 |
| Ale-Ster-64-24 | 1002 |
| Ale-Ster-77-31 | 1006 |
| ÖMö1-7 | 6073 |
| Ängsö-12-402 | 1303 |
| Ängsö-57-419 | 1312 |
| Ängsö-59-422 | 1313 |
| Ängsö-74-430 | 1317 |
| Ängsö-80-432 | 1318 |
| App1-12 | 5830 |
| App1-14 | 5831 |
| App1-16 | 5832 |
| Bag 1 | 9330 |
| Bar 1 | 9332 |
| Bil-3 | 5835 |
| Bön 1 | 9336 |
| Boo2-3 | 5836 |
| Böt 1 | 9339 |
| Böt 4 | 9342 |
| Brösarp-11-135 | 1061 |
| Brösarp-11-138 | 1062 |
| Brösarp-21-140 | 1063 |
| Brösarp-25-142 | 1064 |
| Brösarp-34-145 | 1066 |
| Brösarp-37-149 | 1068 |
| Brösarp-43-152 | 1069 |
| Brösarp-45-153 | 1070 |
| Brösarp-51-157 | 1072 |
| Brösarp-53-159 | 1073 |
| Brösarp-61-162 | 1074 |
| Brösarp-63-163 | 1075 |
| Dja 1 | 9343 |
| **AccessionName** | **AccessionID** |
| Dja 2 | 9344 |
| Djk 3 | 9349 |
| Död 1 | 9351 |
| Död 2 | 9352 |
| Död 3 | 9353 |
| Dör-10 | 5856 |
| Dra1-4 | 5865 |
| Dra2-1 | 5867 |
| Dra-3 | 5860 |
| Dra3-9 | 5870 |
| Eden 15 | 9354 |
| Eden 16 | 9355 |
| Eden 17 | 9356 |
| Eden-1 | 6009 |
| Eden-4 | 8218 |
| Eden-5 | 6010 |
| Eden-6 | 6011 |
| Eden-7 | 6012 |
| Eden-9 | 6013 |
| EdJ 2 | 9363 |
| Eds-9 | 6017 |
| EkN 3 | 9367 |
| EkS 2 | 9369 |
| EkS 3 | 9370 |
| FäL 1 | 9371 |
| Fjä1-2 | 6019 |
| Fjä1-5 | 6020 |
| Fjä2-4 | 6021 |
| Fjä2-6 | 6022 |
| Fly2-1 | 6023 |
| FlyA 3 | 9380 |
| Fri 1 | 9381 |
| Fri 2 | 9382 |
| Fri 3 | 9383 |
| Frö 1 | 9384 |
| Frö 3 | 9385 |
| Gårdby-17-198 | 1132 |
| Gårdby-22-213 | 1137 |
| Gro-3 | 6025 |
| Grön 12 | 9386 |
| Grön 14 | 9388 |
| Grön-5 | 6030 |
| Had 1 | 9390 |
| Had 2 | 9391 |
| Had 3 | 9392 |
| Hag 2 | 9394 |
| Hal 1 | 9395 |
| Ham 1 | 9399 |
| **AccessionName** | **AccessionID** |
| Ham-10-239 | 1366 |
| Ham-13-241 | 1367 |
| Ham-2-2 | 1360 |
| Ham-27-256 | 1374 |
| Ham-6-232 | 1362 |
| Ham-7-233 | 1363 |
| Hel 3 | 9402 |
| Hen-16-268 | 1585 |
| HolA1 1 | 9404 |
| HolA1 2 | 9405 |
| HolA2 2 | 9407 |
| Hov1-10 | 6035 |
| Hov1-7 | 6034 |
| Hov3-2 | 6036 |
| Hov3-5 | 6038 |
| Kal 2 | 9408 |
| Kia 1 | 9409 |
| Kor 1 | 9410 |
| Kor 2 | 9411 |
| Kor 3 | 9412 |
| Kor 4 | 9413 |
| Kru 3 | 9416 |
| Kva 2 | 9418 |
| Lag 1 | 9419 |
| Lan 1 | 9421 |
| Lis-3 | 6041 |
| Löv-1 | 6043 |
| Näs 2 | 9427 |
| Nyl 13 | 9433 |
| Nyl-7 | 6069 |
| Omn-1 | 6070 |
| Omn-5 | 6071 |
| Ost-0 | 8351 |
| Puk 1 | 9436 |
| Puk 2 | 9437 |
| Rev-2 | 6076 |
| Rev-3 | 6077 |
| Röd-17-319 | 1435 |
| Sim 1 | 9442 |
| Sku-30 | 1552 |
| Sparta-1 | 6085 |
| Spro 1 | 9450 |
| Spro 2 | 9451 |
| Spro 3 | 9452 |
| Sr:3 | 6086 |
| Stabby-13 | 1391 |
| Stabby-26 | 1404 |
| Ste 2 | 9453 |
| Ste 3 | 9454 |
| Ste 4 | 9455 |
| Stu-2 | 6087 |
| 1000 | 6090 |
| T1010 | 6091 |
| **AccessionName** | **AccessionID** |
| T1020 | 6092 |
| T1030 | 6093 |
| T1040 | 6094 |
| T1050 | 6095 |
| T1060 | 6096 |
| T1070 | 6097 |
| T1080 | 6098 |
| T1090 | 6099 |
| T1110 | 6100 |
| T1120 | 6101 |
| T1130 | 6102 |
| T1150 | 6103 |
| T1160 | 6104 |
| T450 | 6105 |
| T460 | 6106 |
| T470 | 6107 |
| T480 | 6108 |
| T510 | 6109 |
| T520 | 6110 |
| T530 | 6111 |
| T540 | 6112 |
| T550 | 6113 |
| T570 | 6114 |
| T580 | 6115 |
| T590 | 6116 |
| T610 | 6118 |
| T620 | 6119 |
| T630 | 6120 |
| T640 | 6121 |
| T670 | 6122 |
| T680 | 6123 |
| T690 | 6124 |
| T710 | 6125 |
| T720 | 6126 |
| T730 | 6127 |
| T740 | 6128 |
| T750 | 6129 |
| T760 | 8225 |
| T780 | 6131 |
| T790 | 6132 |
| T800 | 6133 |
| T810 | 6134 |
| T840 | 6136 |
| T850 | 6137 |
| T860 | 6138 |
| T880 | 6140 |
| T890 | 6141 |
| T900 | 6142 |
| T920 | 6144 |
| T930 | 6145 |
| T940 | 6146 |
| T950 | 6147 |
| T960 | 6148 |
| **AccessionName** | **AccessionID** |
| T980 | 6150 |
| T990 | 6151 |
| TÄL 07 | 6180 |
| TÅD 01 | 6169 |
| TÅD 02 | 6170 |
| TÅD 03 | 6171 |
| TÅD 04 | 6172 |
| TÅD 05 | 6173 |
| TÅD 06 | 6174 |
| TAA 03 | 6153 |
| TAA 04 | 6154 |
| TAA 14 | 6163 |
| TAA 17 | 6166 |
| TBÖ 01 | 6184 |
| TDr-1 | 6188 |
| TDr-11 | 6197 |
| TDr-13 | 6198 |
| TDr-14 | 6199 |
| TDr-15 | 6200 |
| TDr-16 | 6201 |
| TDr-17 | 6202 |
| TDr-18 | 6203 |
| TDr-2 | 6189 |
| TDr-22 | 6207 |
| TDr-3 | 6190 |
| TDr-4 | 6191 |
| TDr-5 | 6192 |
| TDr-7 | 6193 |
| TDr-8 | 6194 |
| TDr-9 | 6195 |
| TEDEN 02 | 6209 |
| TEDEN 03 | 6210 |
| TFÄ 04 | 6214 |
| TFÄ 02 | 6212 |
| TFÄ 05 | 6215 |
| TFÄ 06 | 6216 |
| TFÄ 07 | 6217 |
| TFÄ 08 | 6218 |
| TGR 01 | 6220 |
| TGR 02 | 6221 |
| THÖ 03 | 8227 |
| THÖ 08 | 6226 |
| TNY 04 | 6231 |
| TOM 01 | 6235 |
| TOM 02 | 6236 |
| TOM 03 | 6237 |
| TOM 04 | 6238 |
| TOM 06 | 6240 |
| TOM 07 | 6241 |
| Tomegap-2 | 6242 |
| Tos-31-374 | 1247 |
| Tos-75-384 | 1252 |
| **AccessionName** | **AccessionID** |
| Tos-93-391 | 1256 |
| Tos-95-393 | 1257 |
| TRÄ 01 | 6244 |
| Tur 3 | 9469 |
| Tur 4 | 9470 |
| TV-10 | 6258 |
| TV-22 | 6268 |
| TV-30 | 6276 |
| TV-38 | 6284 |
| TV-4 | 6252 |
| TV-7 | 6255 |
| UII2-13 | 8427 |
| UII3-4 | 6413 |
| UIIA 1 | 9471 |
| UIIA 2 | 9472 |
| Ull2-5 | 6974 |
| Vår2-6 | 7517 |
| VårA 1 | 9476 |
| Yst 1 | 9481 |
| Yst 2 | 9482 |
| Fly2-2 | 6024 |
| Stu1-1 | 6088 |
| Vår2-1 | 7516 |
| Hovdala-2 | 6039 |
| Bön2 | ? |
| TGR 02 | 6221 |
| Hovdala-6 | 8307 |
| Bå1-2 | 8256 |
| ÖMö2-1 | 7518 |
| Brö1-6 | 8231 |
| St-0 | 8387 |
| Eden-2 | 6913 |
| Ör-1 | 6074 |
| Fjä1-1 | 8422 |
| Algutstrum | 8230 |
| Gul1-2 | 8234 |
| Tottarp-2 | 6243 |

**Supplementary Table 1 – List of *Arabidopsis thaliana* accessions used in GWA studies**

Accession names and accession IDs for the Arabidopsis lines used in the GWA analysis of All accessions are from Sweden and are a subset of the 1001 genomes project.

The accessions were kindly provided by Caroline Dean at the John Innes Centre, Norwich.
