## Supplementary figures and images for "Variation in the expression of a transmembrane protein influences cell growth in *Arabidopsis thaliana* petals by altering auxin availability"

### Figure S1

#### Slide 1
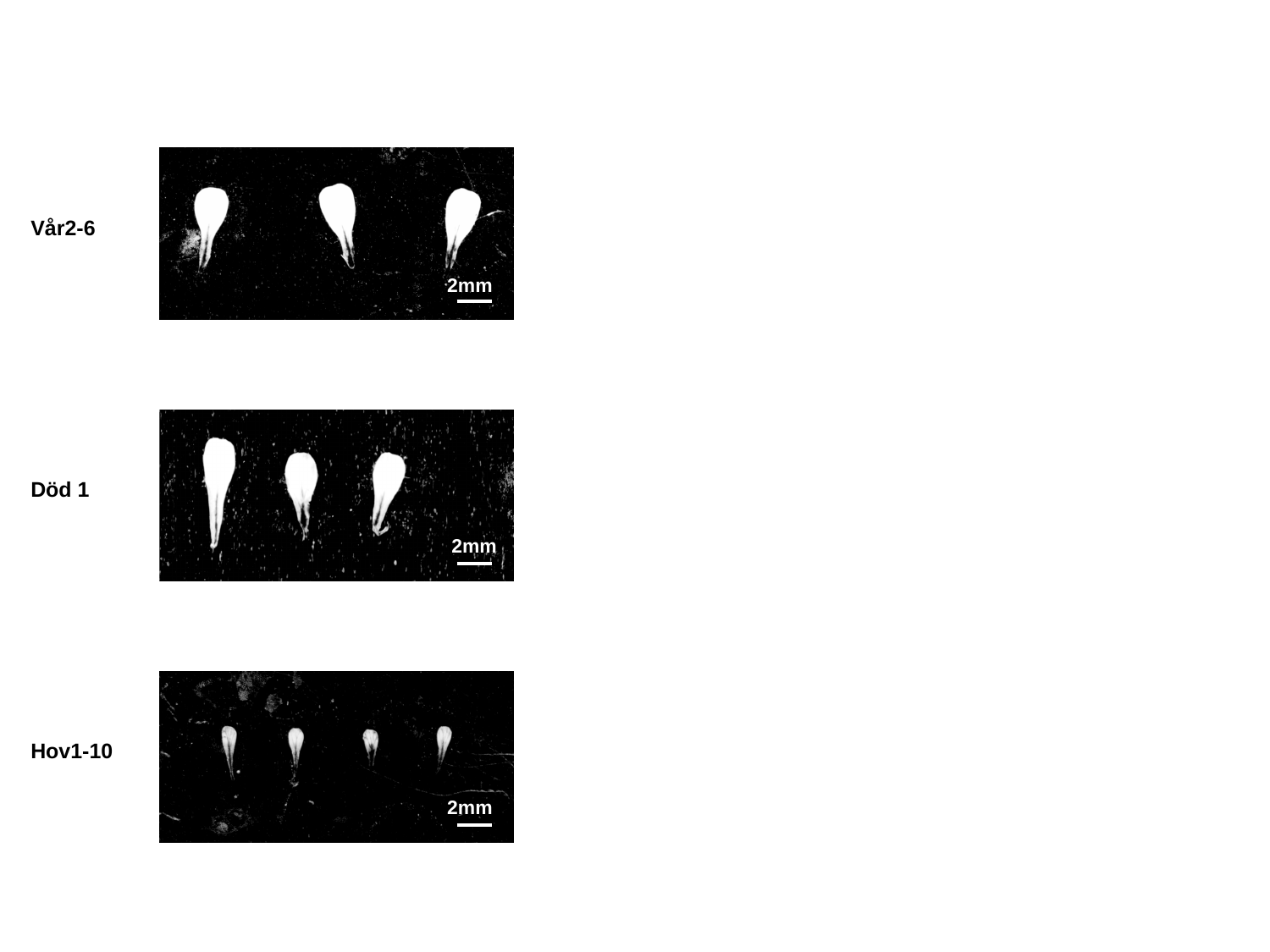

Vår2-6
2mm
Död 1
2mm
Hov1-10
2mm
