## Supplementary material for "Variation in the expression of a transmembrane protein influences cell growth in *Arabidopsis thaliana* petals by altering auxin availability": Figure S2

```
DJA1_AT4G1685   1365 AAATTCCAGGTGTTGGAGATTGTTTTTGATTAAGCTATGGAACCATGGAC   1414
                     |||||||||||||||||||||||||||||||||.||||||||||||||||
TAIR10_AT4G16   1401 AAATTCCAGGTGTTGGAGATTGTTTTTGATTAAACTATGGAACCATGGAC   1450

DJA1_AT4G1685   1415 TTGTCGACTCAGCCACCATCAACAACTGCAATACCATCCTCGAGAATTGC   1464
                     ||||||||||||||||||||||||||||||||||||||||||||||||||
TAIR10_AT4G16   1451 TTGTCGACTCAGCCACCATCAACAACTGCAATACCATCCTCGAGAATTGC   1500

DJA1_AT4G1685   1465 CGTAATAGCTCAGACACCACCACCACCAACAACAACAACAGTGTGGATCG   1514
                     |||||||.|||||      .|||.|.|||||||||||||||||||||||.
TAIR10_AT4G16   1501 CGTAATACCTCAG------TCACTAACAACAACAACAACAGTGTGGATCA   1544

DJA1_AT4G1685   1515 TCCCAGTGACTCAAACACCAACAACAATAACAGTGTGGATCATCCTAATG   1564
                     ||||||||||||||||||||||||||||||||.||||||||||||.||||
TAIR10_AT4G16   1545 TCCCAGTGACTCAAACACCAACAACAATAACATTGTGGATCATCCGAATG   1594

DJA1_AT4G1685   1565 ACATAAACAACAAGAACAATGTTGACAACAAGGACAATAACAGCAGAGAC   1614
                     |||||||.||||||||||||||||||||||||||||||||||||||||||
TAIR10_AT4G16   1595 ACATAAAAAACAAGAACAATGTTGACAACAAGGACAATAACAGCAGAGAC   1644

DJA1_AT4G1685   1615 AAGTAATTAAGTAGGAAAAATCT-CGGTTTAGATGATACCG----ATCGG   1659
                     ||||||||||.|||||||.| || |||||||||||||||||    |||||
TAIR10_AT4G16   1645 AAGTAATTAAATAGGAAACA-CTCCGGTTTAGATGATACCGATCTATCGG   1693
			VRN2 term.
DJA1_AT4G1685   1660 ATTGTAACTTATTCTTCTTTCTTAAAAAAATTGTTTAGGAGCAAACGAA-   1708
                     ||||||||||||||||||||||||||||||||||||||||||||||.||
TAIR10_AT4G16   1694 ATTGTAACTTATTCTTCTTTCTTAAAAAAATTGTTTAGGAGCAAACAAAG   1743

DJA1_AT4G1685   1709 ATTTTAATTTGTTAGTGTGTATTCAACTGATTACATTTTTAGTT--GAAA   1756
                     ||||| ||||||||  ||||||||||||||||||||||||||||  .|||
TAIR10_AT4G16   1744 ATTTT-ATTTGTTA--GTGTATTCAACTGATTACATTTTTAGTTAAAAAA   1790

DJA1_AT4G1685   1757 ATGAATCCTGTTTAATTATTATTTTTTAATTAAAAACTGAAAAATTAAGA   1806
                     |||.||.||..||||           |||.||||.||||||||| |||||
TAIR10_AT4G16   1791 ATGGATTCTCCTTAA-----------TAACTAAAGACTGAAAAA-TAAGA   1828

DJA1_AT4G1685   1807 AAAGTTTACTTAGTTTTTCTTTATGACTTGAGAAAAAGCT--------CC   1848
                     .||||||.||||.|||||||||.|||||||||||||||||        ||
TAIR10_AT4G16   1829 TAAGTTTCCTTAATTTTTCTTTTTGACTTGAGAAAAAGCTCCTCTAGACC   1878

DJA1_AT4G1685   1849 TCTCGTCAATAGGAGTTATATATAGATCAACTACATAACATAAATATATA   1898
                     |||.||.||||||||||||||||..|||||.||||||||||||
TAIR10_AT4G16   1879 TCTAGTAAATAGGAGTTATATATTAATCAAGTACATAACATAA-------   1921

DJA1_AT4G1685   1899 TATATATATATATATATATATATATATATATATATTAAGTGCAAATAGAT   1948
                                            |.|||||||||||||||||||||||||
TAIR10_AT4G16   1922 -----------------------AAATATATATATTAAGTGCAAATAGAT   1948

DJA1_AT4G1685   1949 TGAAAACAAATCAAGAAATTAATTAAGACAGAGTGATTAAGCTTAAAACC   1998
                     ||||||||||||||||||||||||||||||.|||||||||||||||||||
TAIR10_AT4G16   1949 TGAAAACAAATCAAGAAATTAATTAAGACACAGTGATTAAGCTTAAAACC   1998
			KSK txn start site
DJA1_AT4G1685   1999 CC   2000
                     ||
TAIR10_AT4G16   1999 CC   2000
```

**Supplementary Figure 2.** Alignment (EMBOSS Needle) of *KSK* sequences upstream of the transcription start site in ecotypes Col-0 and Dju-1.
